# Absence of severe COVID-19 in patients with clonal mast cells activation disorders: effective anti-SARS-CoV-2 immune response

**DOI:** 10.1101/2021.09.01.458516

**Authors:** Julien Rossignol, Amani Ouedrani, Cristina Bulai Livideanu, Stéphane Barete, Louis Terriou, David Launay, Richard Lemal, Celine Greco, Laurent Frenzel, Cecile Meni, Christine Bodemere-Skandalis, Laura Polivka, Anne-Florence Collange, Hassiba Hachichi, Sonia Bouzourine, Djazira Nait Messaoud, Mathilde Negretto, Laurence Vendrame, Marguerite Jambou, Marie Gousseff, Stéphane Durupt, Jean-Christophe Lega, Jean-Marc Durand, Caroline Gaudy, Gandhi Damaj, Marie-Pierre Gourin, Mohamed Hamidou, Laurence Bouillet, Edwige Le Mouel, Alexandre Maria, Patricia Zunic, Quentin Cabrera, Denis Vincent, Christian Lavigne, Etienne Riviere, Clement Gourguechon, Anne Brignier, Ludovic Lhermitte, Thierry Jo Molina, Julie Bruneau, Julie Agopian, Patrice Dubreuil, Dana Ranta, Alexandre Mania, Michel Arock, Isabelle Staropoli, Olivier Tournilhac, Olivier Lortholary, Olivier Schwartz, Lucienne Chatenoud, Olivier Hermine

## Abstract

Mast cells are key actors of innate immunity and Th2 adaptive immune response which counterbalance Th1 response, critical for anti-viral immunity. Clonal Mast Cells Activation Disorders (cMCADs) such as mastocytosis and clonal mast cells activation syndrome are characterized by an abnormal mast cells accumulation and/or activation. No data have been published on the anti-viral immune response of patients with cMCADs. The aims of the study were to collected, in a comprehensive way, outcomes of cMCADs patients who experienced a biologically-proven COVID-19 and to characterize both anti-endemic coronaviruses and specific anti-SARS-CoV-2 immune responses in these patients. Clinical follow-up and outcome data were collected prospectively for one year within the French rare disease network CEREMAST encompassing patients from all over the country. Anti-SARS-CoV-2 and anti-endemic coronaviruses specific T-cells were assessed with an enzyme-linked immunospot assay (EliSpot) and anti-SARS-CoV-2 humoral response with dosage of circulating levels of specific IgG, IgA and neutralizing antibodies. Overall, 32 cMCADs patients were identified. None of them required non-invasive or mechanical ventilation; two patients were hospitalized to receive oxygen and steroid therapy. In 21 patients, a characterization of the SARS-CoV-2-specific immune response has been performed. A majority of patients showed a high proportion of circulating SARS-CoV-2-specific interferon (IFN)-γ producing T-cells and high levels of anti-Spike IgG antibodies with neutralizing activity. In addition, no defects in anti-endemic coronaviruses responses were found in patients with cMCADs compared to non-cMCADs controls. Patients with cMCADs frequently showed a spontaneous IFN-γ T-cell production in absence of any stimulation that correlated with circulating basal tryptase levels, a marker of mast cells burden. These findings underscore that patients with cMCADs might be not at risk of severe COVID-19 and the spontaneous IFN-γ production might explain this observation.

**Author Summary:** Mast cells are immune cells involved in many biological processes including the anti-microbial response. However, previous studies suggest that mast cells may have a detrimental role in the response against viruses such as SARS-CoV-2, responsible for COVID-19. When a mutation occurs in mast cells, it can lead to a group of diseases called clonal mast cells activation disorders (cMCADs), characterized by deregulated activation of these cells. Hence, patients with cMCADs might be more susceptible to severe COVID-19 than general population.

We therefore conducted a 1-year study in France to collect data from all cMCADs patients included in the CEREMAST rare disease French network and who experienced COVID-19. Interestingly, we did not find any severe COVID-19 (i.e. requiring non-invasive or mechanical ventilation) in spite of well-known risk factors for severe COVID-19 in a part of cMCADs patients.

We then have studied the immune response against SARS-CoV-2 and other endemic coronaviruses in these patients. We did not observe any abnormalities in the immune response either at the level of T and B lymphocytes. These findings underscore that these patients might not be at risk of severe COVID-19 as one might have feared.

## Introduction

Clonal mast cells activation disorders (cMCADs) are a spectrum of heterogeneous diseases ranging from monoclonal mast cells activation syndrome (MMAS) to mastocytosis characterized by the activation and/or accumulation of pathological mast cells(1). In adults, the most frequent form of cMCADs is indolent systemic mastocytosis (ISM). Advanced mastocytosis (including aggressive systemic mastocytosis, mast cells leukemia, and systemic mastocytosis with an associated hematological neoplasm) are rarer and linked to poor prognosis(2,3).

COVID-19 is a potentially fatal infectious disease caused by the emerging SARS-CoV-2 virus, which has caused a global pandemic(4). At the pathophysiological level, there is compelling data to support the major role of interferons (IFNs) in the control of disease. It includes type I IFN, produced by plasmacytoid dendritic cells, and IFN-γ (type III IFN), produced by adaptive T-cells in the early and later phases of the disease respectively(5–8).

Well-established capacity of mast cells to drive Th2 responses(9,10), which counterbalance Th1 responses, could make one fears that it impairs anti-viral immunity in patients with cMCADs. In addition, *in vitro* studies found that histamine blocks the activity of human plasmacytoid dendritic cells, thereby further impacting on anti-viral responses(11). Furthermore, mast cells may contribute to COVID-19-induced inflammation by releasing pro-inflammatory cytokines such as interleukin (IL-)1, IL-6 and tumor necrosis factor (TNF) and may also exacerbate the lung lesions *via* degranulation(12,13). Hence, patients with cMCADs could have been more susceptible to severe COVID-19.

Over one year, we prospectively collected data from all patients with cMCADs (MMAS and mastocytosis) included in the CEREMAST rare disease French network and who experienced a biologically proven COVID-19. Here we aimed at describing the clinical course, outcome and immunological characteristics of those patients.

COVID-19 was diagnosed in presence of a positive SARS-CoV-2 PCR on nasal swab or of COVID-19 suggestive symptoms and positive anti-SARS-CoV-2 serology. Patients with COVID-19 suggestive symptoms without biological evidence of SARS-CoV-2 infection were excluded from the study. First, all patients included in the CEREMAST network were sent a request to report any COVID-19 episode. If they reported one, the specialist physician in charge of the patient subsequently confirmed the case. Second, all specialists in the CEREMAST network were contacted to report additional cases not reported by patients themselves. Eventually, interrogation of the computerized registry of the Paris Public Hospitals Public Assistance (Assistance Publique des Hôpitaux de Paris (APHP)) searched for additional inpatient cases.

To assess the SARS-CoV-2-specific T and B lymphocyte responses, we respectively analysed specific T-cells reactivities using an enzyme-linked immunospot assay (EliSpot), and circulating levels of specific IgG, IgA and neutralizing antibodies, and compared them to non cMCADs controls. The EliSpot measured IFN-γ production after a short stimulation of freshly isolated PBMC with different pools of peptide derived from a scan through five SARS-CoV-2 proteins. Six pools were tested: S1 for Spike glycoprotein N-terminal fragment, S2 for Spike glycoprotein C-terminal fragment, M for Membrane protein, N for Nucleoprotein, E for Envelope small membrane protein and AP3a for ORF3a protein. T-cells response was also tested in 32 COVID-19 positive controls with mild-moderate (n=17) to severe (n=15) forms.

## Results and discussion

### Characteristics and outcomes of cMCAD patients with COVID-19

From February 1rst 2020 to February 1rst 2021, 32 patients with cMCADs and COVID-19 were prospectively identified by the CEREMAST network (Figure 1). Among them, 18 patients were initially identified with the questionnaire, 14 patients were secondarily identified directly by the referring physician for cMCADs, and no additional inpatients cases were retrieved in the APHP database.

**Figure 1:**
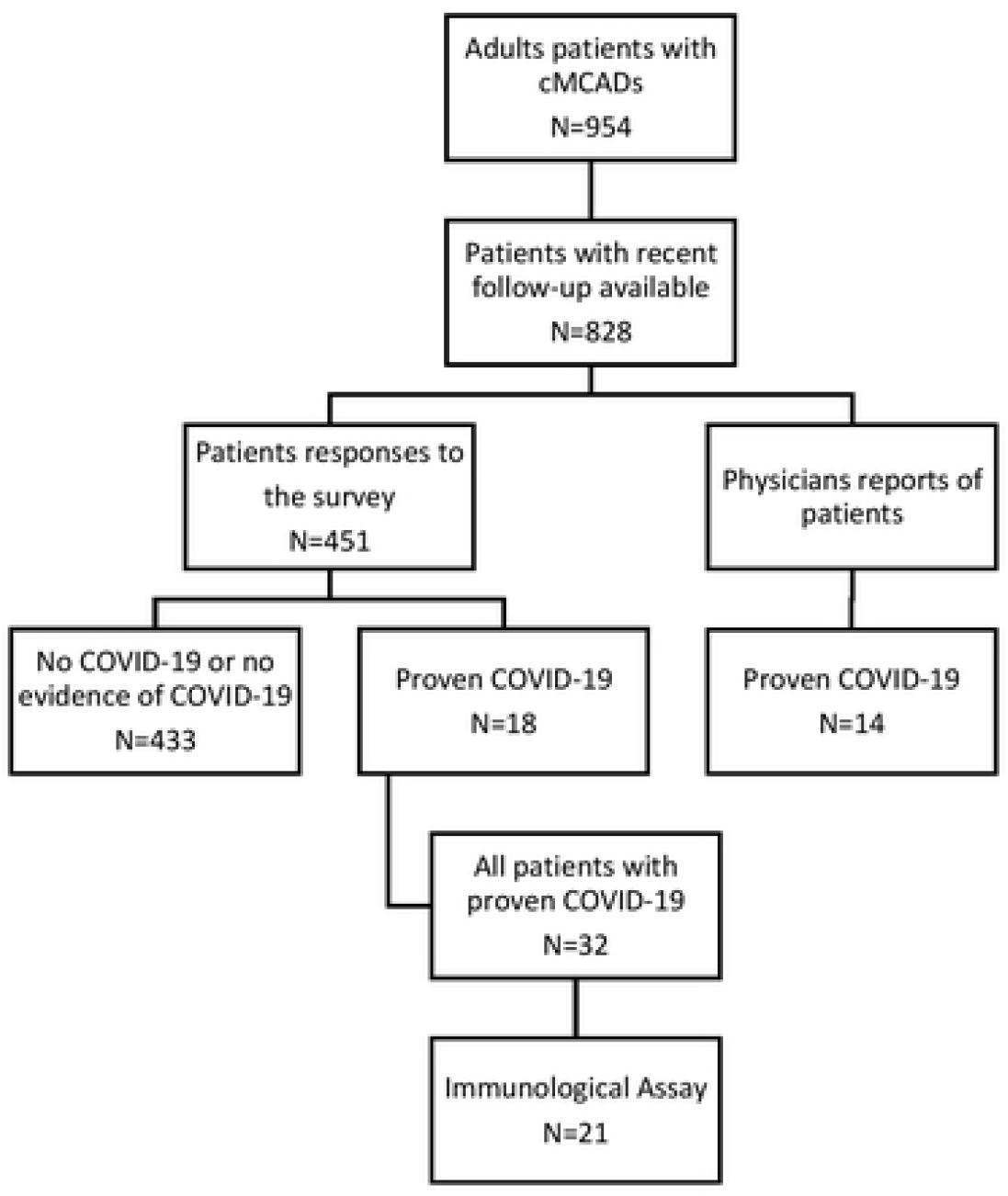
Flowchart of patients with cMCADs and proven COVID-19 identification. cMCADs: clonal mast cells activation disorders.

Characteristics and outcomes of patients with cMCADs and COVID-19 are detailed in Table 1. Patients were predominantly females (59.4%) with a median age of 49.7 years (ranging from 25.6 to 76.4 years). The subtype of cMCADs among the 32 patients was cutaneous mastocytosis or mastocytosis in the skin in 14 patients, ISM in 15 patients, one patient with smoldering systemic mastocytosis (SSM) and 2 patients with MMAS. Among 21 patients in whom a genetic analysis was performed 18 (85.7%) were carriers of the D816V *KIT* mutation. Ten patients (31.3%) had a history of severe anaphylactic reaction and median basal serum tryptase before any clinical or biological sign of COVID-19 was 13.0 μg/L (ranging from 2.7 to 163.0 μg/L). Risk factors predisposing to severe COVID-19 were present in 13/32 patients (40.6%) and 4 of them had at least 2 risk factors(14). These risk factors were BMI > 30 (N=4), age > 65 years (N= 4), current cytoreductive therapy (midostaurin) or recent (<1 year) administration of cladribine (2CDA) (N=3), cardiovascular condition including arterial hypertension and chronic heart failure (N=7) and diabetes (N=2). At the time of the SARS-CoV-2 infection, 23/32 (71.9%) patients were receiving symptomatic treatments (anti-H1 and/or anti-H2 and/or montelukast), one patient with SSM was receiving midostaurin after failure of 2CDA and one patient with ISM had recently received 2CDA.

**Table 1:** Characteristics and outcomes of patients with cMCADs and COVID-19. Patients are classified according to their age. #: Patient number. F: female. M: Male. Age (Years). MIS: Mastocytosis in the Skin. CM: Cutaneous Mastocytosis. MMAS: Monoclonal mast cells activation syndrome. SSM: Smoldering systemic mastocytosis. Risk factors: Risk factors for severe COVID-19(14). NA: not available. BMI: Body mass index. Cytoreductive therapy: midostaurin or current/recent (< 1 year) administration of 2CDA. COVID-19 scale: WHO COVID-19 clinical progression scale (15). *Patients treated with corticosteroid therapy.

| # | Sex | Age | BMI | cMCADs | *KIT* mutation | History of severe anaphylaxis | Risk factors | Tryptase µg/L | Treatment: anti-H1 | Treatment: anti-H2 | Treatment: montelukast | Cytoreductive therapy | COVID-19 scale | Symptoms of cMCADs | Fever | Anosmia/Ageusia | PCR SARS-CoV-2 | Serology SARS-CoV-2 |
| --- | --- | --- | --- | --- | --- | --- | --- | --- | --- | --- | --- | --- | --- | --- | --- | --- | --- | --- |
| 1 | F | 25.6 | 17.4 | ISM | WT | no | no | 3.9 | no | yes | no | no | 2 | Unchanged | yes | yes | yes | Positive |
| 2 | F | 26.7 | 24.3 | CM | D816V | no | no | 4.4 | yes | no | yes | no | 2 | Increased | yes | yes | yes | Negative |
| 3 | F | 29.9 | 23.9 | MIS | NA | no | no | 4.7 | no | no | no | no | 2 | Unchanged | no | no | no | Positive |
| 4 | F | 33.4 | 28.1 | CM | D816V | yes | no | 6.6 | yes | no | no | no | 2 | Unchanged | yes | yes | no | Positive |
| 5 | F | 35.5 | 33.1 | MIS | NA | no | yes | 7.4 | no | no | no | no | 2 | Unchanged | no | yes | yes | Positive |
| 6 | F | 38.3 | 18.4 | ISM | D816V | yes | no | 7.0 | no | no | no | no | 2 | Increased | no | no | yes | Negative |
| 7 | F | 40.3 | 21.0 | CM | WT | no | no | 8.1 | yes | no | no | no | 2 | Unchanged | no | yes | no | Positive |
| 8 | F | 41.4 | 22.5 | ISM | NA | no | no | 7.7 | yes | no | no | no | 2 | Decreased | yes | yes | yes | Positive |
| 9 | F | 42.3 | 20.0 | MMAS | D816V | yes | no | 18.5 | yes | no | yes | no | 2 | Increased | no | yes | yes | Positive |
| 10 | M | 42.4 | 33.5 | ISM | D816V | no | yes | 2.7 | yes | no | no | no | 2 | Decreased | yes | yes | yes | Positive |
| 11 | F | 43.3 | 25.0 | CM | WT | yes | no | 13.0 | yes | no | yes | no | 2 | Unchanged | yes | no | yes | Positive |
| 12 | F | 43.3 | 24.3 | ISM | D816V | yes | yes | 60.0 | yes | yes | no | yes | 2 | Increased | yes | yes | yes | Positive |
| 13 | F | 43.8 | 25.1 | MIS | NA | no | no | 13.0 | no | no | no | no | 2 | Unchanged | yes | yes | yes | Positive |
| 14 | M | 45.3 | 30.8 | MMAS | D816V | yes | yes | 45.0 | yes | no | no | no | 2 | Unchanged | yes | no | no | Positive |
| 15 | F | 48.1 | 17.6 | ISM | D816V | yes | no | 99.8 | yes | no | yes | no | 2 | Unchanged | no | yes | yes | Positive |
| 16 | M | 49.1 | 27.4 | ISM | D816V | no | no | 14.8 | yes | yes | no | no | 2 | Increased | yes | yes | no | Positive |
| 17 | M | 50.3 | 23.2 | MIS | NA | no | no | 18.6 | yes | no | no | no | 2 | Unchanged | no | no | yes | Positive |
| 18 | M | 51.7 | 26.4 | ISM | D816V | no | no | 42.8 | yes | no | no | no | 2 | Unchanged | yes | yes | yes | Positive |
| 19 | F | 52.2 | 19.5 | ISM | NA | no | no | 7.9 | yes | no | yes | no | 2 | Unchanged | yes | no | no | Positive |
| 20 | F | 52.4 | 30.1 | CM | D816V | no | yes | 37.2 | yes | no | no | no | 5* | Increased | yes | no | yes | Positive |
| 21 | F | 52.9 | 17.5 | ISM | D816V | no | yes | 38.6 | yes | no | no | no | 2 | Unchanged | yes | yes | yes | Negative |
| 22 | F | 53.1 | 27.0 | MIS | NA | no | no | 31.4 | no | no | no | no | 2 | Unchanged | yes | yes | yes | Positive |
| 23 | M | 56.0 | 26.8 | ISM | NA | no | yes | 19.0 | no | no | no | no | 2 | Unchanged | yes | yes | no | Positive |
| 24 | F | 59.6 | 21.6 | MIS | NA | no | no | 56.0 | no | no | no | no | 2 | Increased | no | no | yes | Positive |
| 25 | M | 60.7 | 26.6 | CM | D816V | no | no | 12.0 | no | no | no | no | 1 | Unchanged | no | no | yes | Positive |
| 26 | M | 62.2 | 26.5 | ISM | D816V | no | yes | 6.1 | yes | no | no | no | 2 | Unchanged | yes | no | no | Positive |
| 27 | M | 62.2 | 24.2 | CM | D816V | no | no | 18.6 | yes | no | no | no | 2 | Unchanged | yes | yes | yes | Positive |
| 28 | M | 63.2 | 28.1 | MIS | NA | yes | yes | 11.2 | yes | no | yes | no | 2 | Unchanged | no | yes | yes | Positive |
| 29 | M | 65.6 | 26.7 | ISM | D816V | yes | yes | 27.8 | no | no | no | no | 5* | Unchanged | yes | no | yes | Positive |
| 30 | F | 73.9 | 21.2 | SSM | D816V | no | yes | 163.0 | no | yes | no | yes | 2 | Unchanged | yes | no | no | Positive |
| 31 | M | 76.2 | 25.4 | ISM | D816V | yes | yes | 8.5 | yes | no | no | no | 3 | Decreased | yes | no | yes | Positive |
| 32 | M | 76.5 | 27.8 | ISM | NA | no | yes | 7.6 | no | yes | no | no | 2 | Increased | yes | no | yes | Positive |

Regarding the diagnosis of COVID-19, 23/32 (71.9%) patients had a positive SARS-CoV-2 PCR on nasal swab, and 9/32 had either a negative or did not underwent SARS-CoV-2 PCR on nasal swab due to a non-availability of the procedure at the time of infection but had a positive serology. A large majority of patients (29/32) seroconverted during their follow up. As expected, patients with subsequent sera available (N=4) had all negated their serology after a median follow up of 33.0 weeks. Regarding the symptoms of COVID-19, fever (>38°C) was found in 22/32 (68.8%) patients and anosmia and/or ageusia in 18/32 (56.3%) patients. Frostbite of the toes persisting after infection was found in one patient without risk factors. Only two patients required hospitalization for corticosteroid therapy and oxygen therapy (stage 5 according to WHO COVID-19 clinical progression scale(15)) but none required non-invasive or mechanical ventilation. Interestingly, 8/32 (25.0%) of patients reported an increase in signs of mast cells activation during the COVID-19 while 3/32 (9.4%) reported a decrease. No recurrence of the infection has been reported.

Overall, no severe COVID-19 disease case was observed in this comprehensive series of patients despite the high prevalence of risk factors (obesity, advanced age, cardiovascular conditions or immunosuppressive treatments). This finding confirms the recently published data from an international study(16). However, our study extends our knowledge on cMCADs and COVID-19 due to the exhaustive nature of the inclusion that concerned the CEREMAST rare disease network, which encompasses cMCADs in the entire French population. Indeed, when a patient with mastocytosis not referenced in the network was hospitalized for COVID-19 disease in an intensive care unit, the local or national reference centers were systematically contacted to obtain an expert opinion on potential drug contraindications due to the mandatory precautions needed for anaesthesia. The exhaustivity of the recruitment of patients with advanced mastocytosis and severe or critical COVID-19 was confirmed through consultation of the computerized registry of APHP that did not retrieve any inpatient unknown to the CEREMAST network. For obvious reasons, the only bias is that we cannot be fully exhaustive concerning patients with asymptomatic, mild or moderate forms of COVID-19 disease that did not require hospitalization nor special advice from their referent physicians.

### Characterization of anti-SARS-CoV-2 and anti-endemic coronaviruses specific T-cells with an enzyme-linked immunospot assay in patients with cMCADs

The anti-SARS-CoV-2 specific cellular and humoral immune responses were studied in 21 cMCADs within a median of 24 weeks [IQR 7-36] from infection.

Overall, 20/21 cMCADs patients have developed a specific T-cell response against at least one of SARS-CoV-2 peptide pools tested. Relatively modest intensities were found. Median intensities for S1 pool were 37 SFU/10^3^ CD3 [IQR 24-130], S2 pool 108 SFU/10^3^ CD3 [IQR 23-201], M pool 62 SFU/10^3^ CD3 [IQR 21-146], N pool 78 SFU/10^3^ CD3 [IQR 48-256] and AP3a pool 23 SFU/10^3^ CD3 [IQR 10-66]. The only patient who did not develop any specific T-cell response (#6), was young (38 years old), diagnosed with PCR on nasal swab, has presented a mild COVID-19 and had no history of immune deficiency or immunosuppressive therapy.

Interestingly, cMCADs patients developed similar reactivity to those found in the control group with mild-moderate COVID-19 in terms of frequency and intensity for S2, M, N and AP3a pools (Figure 2). However, response was significantly lower for Spike glycoprotein N-terminal fragment pool in cMCADs Vs COVID-19 mild-moderate controls (37 SFU/10^3^ CD3 [IQR 24-130] Vs 114 SFU/10^3^ CD3 [IQR 52-289] respectively, p=0.0288). Similarly, when compared to severe COVID-19 controls, we found significantly lower SARS-CoV-2 specific T-cell response in cMCADs patients (p<0.001) except for N pool (Figure 2). Of note, in all groups we did not detect (or very few) specific T-cells for SARS-CoV-2 envelope small membrane protein. Anti-SARS-CoV-2 immune profiles of patient #20 with grade 5 WHO COVID-19 clinical progression scale as well as patient #30 who have received recent administration of 2CDA did not seem different from others patients.

**Figure 2:**
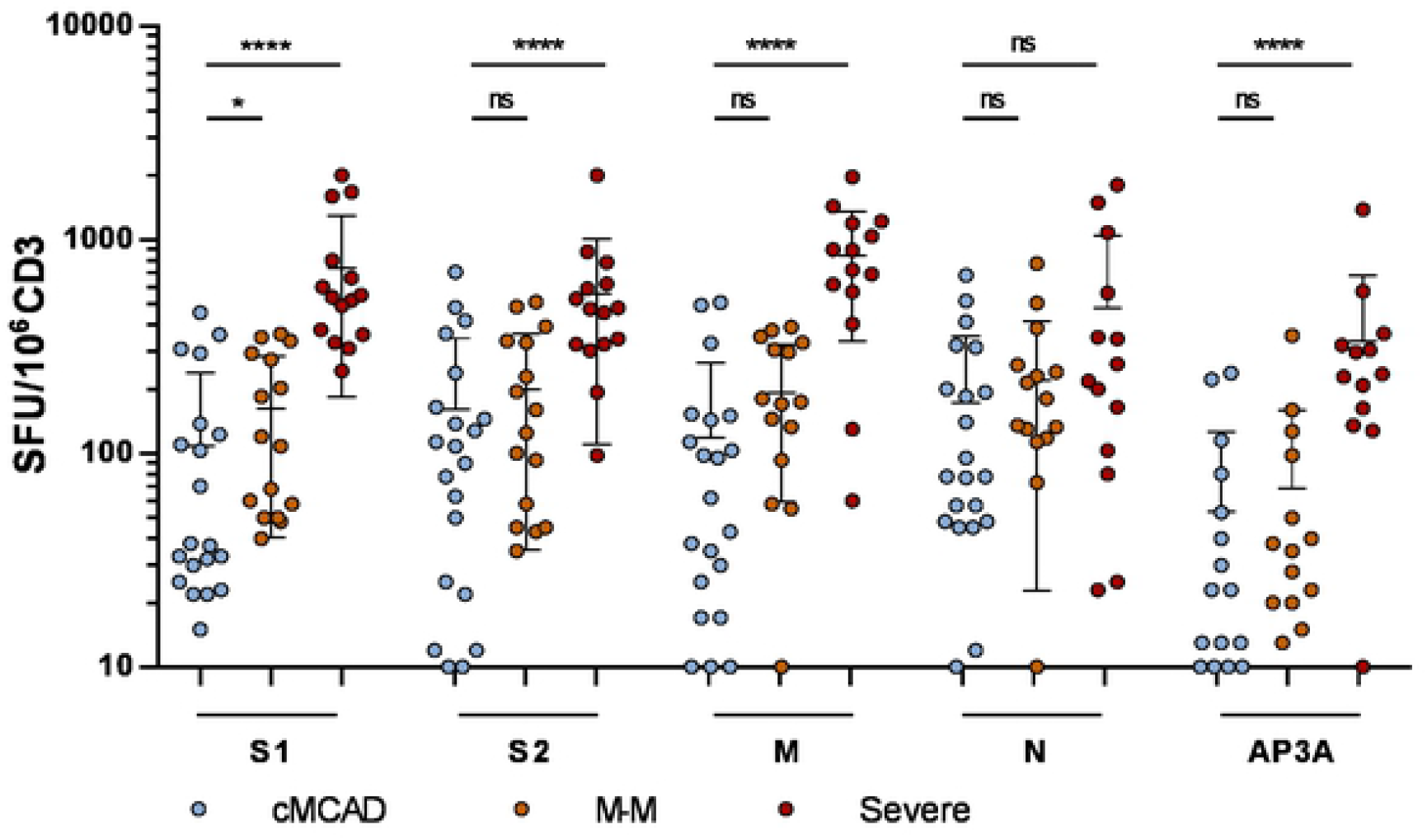
Quantification of SARS-CoV-2 specific T-cells responses using EliSpot. Results were expressed as spot forming unit (SFU)/10^6^ CD3^+^ T-cells after subtraction of background values from wells with non-stimulated cells. Negative controls were PBMC in culture medium. Positive controls were PHA-P and CEFX Ultra SuperStim Pool. SARS-Cov-2 peptide pools tested were derived from a peptide scan through SARS-CoV-2 Spike glycoprotein (S1: N-terminal fragment, S2: C-terminal fragment), Membrane protein (M), Nucleoprotein (N), and ORF3a protein (AP3a). cMCADs: convalescent patients with clonal Mast Cells Activation Disorders. M-M: Convalescent controls with mild to moderate COVID-19 forms. Severe: Convalescent controls with severe COVID-19 forms. NS: non-significant; *, P < 0.05; ****, P < 0.0001.

To evaluate the global anti-coronavirus immune response in patients with cMCADs, we studied T-cell specific response against Spike glycoprotein of Human alpha and beta-coronavirus HCoV-229E, HCoV-NL63, HCoV-OC43 and HCoV-HKU1. Two pools of peptide were tested (S1 and S2) as for SARS-CoV-2 Spike glycoprotein (S1 figure). No significant differences were found when comparing response in cMCADs patients and non-cMCADs controls. Our study did not find any defects in anti-endemic coronaviruses responses in patients with cMCADs with comparable reactivities in terms of frequency and intensity compared to non-cMCADs controls. Same observation was found when comparing response in cMCADs and controls to the EliSpot positive control CEFX Ultra SuperStim Pool containing 176 known peptide epitopes derived from a broad range infectious agent: the IFN-γ production was similar in mastocytosis as in non-mastocytosis patients (S2 figure).

### Characterization of anti-SARS-CoV-2 humoral response in patients with cMCADs

In parallel with EliSpot, SARS-CoV-2 specific IgG and IgA antibodies were studied with a very sensitive technique: The S-flow assay in 15 cMCADs patients. Fourteen of 15 were positive for IgG and 7 of 15 for IgA. The IgG negative patient was the one with negative EliSpot (#6). A viral pseudo-particle neutralization assay was used to determine if IgG were neutralizing. In 12/14 (86%) of IgG seropositive patients we detected neutralizing antibodies. We report here a high prevalence of anti-SARS-CoV-2 seropositivity with high titter of neutralizing antibodies (S3 figure).

Taking all these observations into account, it is believed that patients with mastocytosis were able to develop an effective and protective Th1 cell response against SARS-CoV-2 contrary to what initially expected. In fact, as numerous studies have reported a major role of mast cells in the Th2 immune polarization, it was thought that patients with cMCADs were at high risk for severe COVID-19 with potential lack in Th1 cell anti-viral response. Strikingly, we observed no severe infections in our patients, and only two patients with 1 and 3 risk factors respectively had to be hospitalized for low-flow oxygen therapy with favourable outcome. Almost all our patients developed a SARS-CoV-2 specific T-cell response with equivalent reactivities in term of frequency to non-cMCADs controls. Thus, mast cells from patients with cMCADs do not appear to elicit a worse Th1 response. However, as lower intensities against SARS-CoV-2 Spike glycoprotein were observed in cMCADs, we cannot exclude an impact of mast cells on the amplitude of the Th1 response, and it raises concerns about post-immunisation cellular response. Besides recent works have shown that mast cells may also play a role in the Th1 response, especially in anti-viral response(17–19). Mast cell could play a role in the balance Th1/Th2 potentially important for preventing severe forms of COVID-19.

### IFN-γ spontaneous production in EliSpot assays of patients with cMCADs

Reading EliSpot’s plates revealed an interesting observation: significantly higher backgrounds were found in cMCADs when comparing with non-cMCADs control group. In non-stimulated wells, containing PBMC in culture medium without any peptide pool, we accounted more than 10 small spots/2 10^5^ CD3+ in 10/24 cMCADs patients (with history or not of COVID-19) versus 3/31 non-cMCADs controls (Fisher’s Exact Test: p=0,009) and 2/11 in controls patients with idiopathic mast cell activation syndrome (Figure 3A).

**Figure 3:**
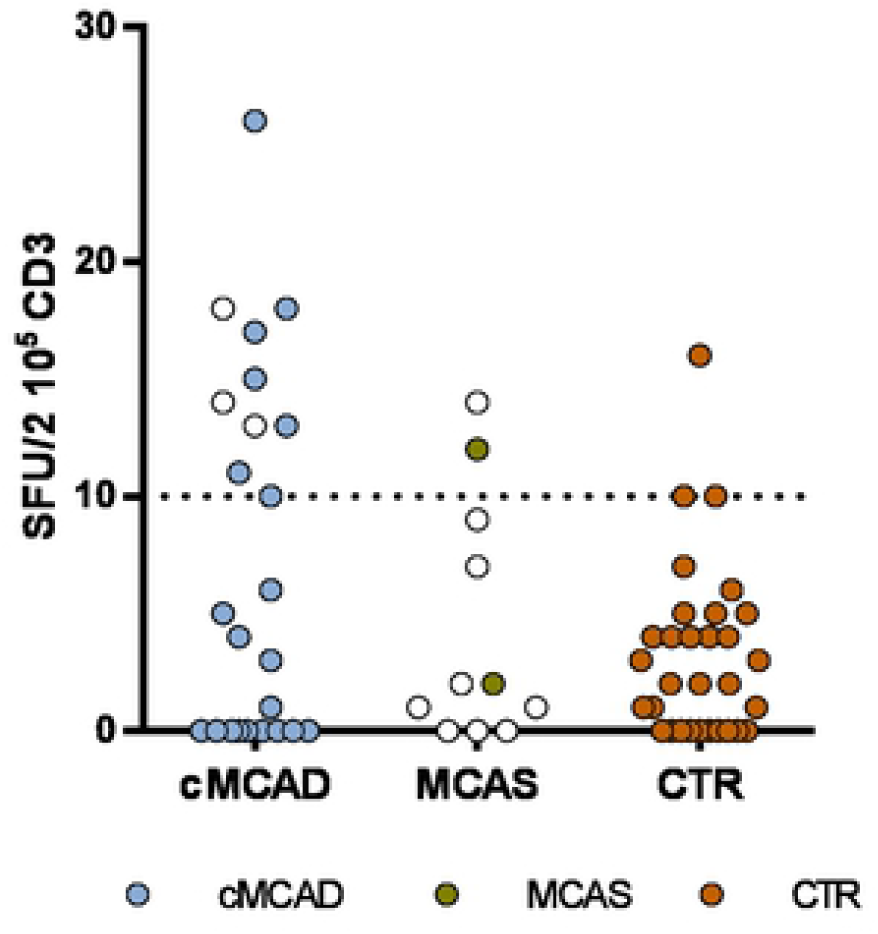
IFN-γ spontaneous production in EliSpot assays of patients. **A.** cMCADs: patients with clonal Mast Cells Activation Disorders. MCAS: patients with idiopathic mast cell activation syndrome. CTR: convalescent controls without cMCADs or MCAS. Empty circle: no COVID-19. Filled circle: history of COVID-19. **Pictures of EliSpot assays. B:** Well with non-stimulated PBMC from COVID-19 control without cMCADs. **C**: Well with non-stimulated PBMC from COVID-19 cMCADs patient. **D**: Well with PBMC from COVID-19 control without cMCADs after stimulation for 18-20h using individual 15-mers 11-aa overlapping peptide pools derived from SARS-CoV-2 N-terminal fragment Spike protein. **E**: Well with PBMC from COVID-19 cMCADs patient after stimulation for 18-20h using individual 15-mers 11-aa overlapping peptide pools derived from SARS-CoV-2 N-terminal fragment Spike protein.

Of note, size and intensity of SARS-CoV-2 specific spots were much greater than background spots (Figure 3B-E). Thus, adjusting settings of EliSpot Reader made it possible to count SARS-CoV-2 specific spots accurately and objectively.

The phenomenon observed resulted from a spontaneous IFN-γ release in the absence of any stimulation. As we tested total PBMC, our assay did not allow to identify the specific IFN-γ producing population. PBMC include T and B-cells, natural killer (NK) cells, monocytes and other myeloid cells such as dendritic cells. The spontaneous IFN-γ release could be associated with elevated levels of basal T-cell activation. Liu et al. (20) reported an increase of activated CD4+ T-cells (CD4+CD38+HLA-DR+) and CD8+ T-cells (CD8+CD38+HLA-DR+) frequencies in individuals with high background compared to those with low background in HIV-1-seronegative individuals. Another hypothesis would involve NK cells in maintaining an elevated baseline IFN-γ level. NK cells of polyallergic patients spontaneously released higher amounts of IFN-γ, interleukin (IL)-4, IL-5 and IL-13 compared to healthy individuals (21) demonstrating an *in vivo* activation of NK cells in atopic patients and suggesting that NK cell might be involved in unbalanced cytokine network in allergic inflammation.

Accordingly, we aimed to understand whether spontaneous IFN-γ release in cMCADs patients was related to NK cells similarly to what observed in polyallergic patients. Human NK cells can be divided into two subsets NK1 and NK2 based on their capacity to secrete IFN-γ (22). NK1 secrete IFN-γ and inhibits IgE synthesis in allergy (21). Level of total IgE could therefore be used as indirect marker of NK1 activation. Thus, we determined total IgE in 17 cMCADs sera. Low IgE levels were found in our cohort, the median value was 20 UI/ml [IQR 12.8 – 36.5] and no correlation between total IgE levels and spontaneous IFN-γ release was found. Further investigations are needed to characterize NK cells in patients with mastocytosis.

We aimed then to correlate with patient characteristics. No correlation was found with age, current symptomatic treatments, history of anaphylaxis or the presence of *KIT* D816V mutation. Interestingly, we found that basal serum tryptase level was correlated with spontaneous IFNγ release in patients with CM, MIS and ISM (Figure 4, R2=0.61, p<0.0001). Although it seems very unlikely that tryptase is directly involved in this phenotype (especially since patients with advanced mastocytosis have very high tryptase level without any known protection against infection) we believe that it reflects a link between clonal mast cells burden and IFN-γ release in patient with non-advanced mastocytosis.

**Figure 4:**
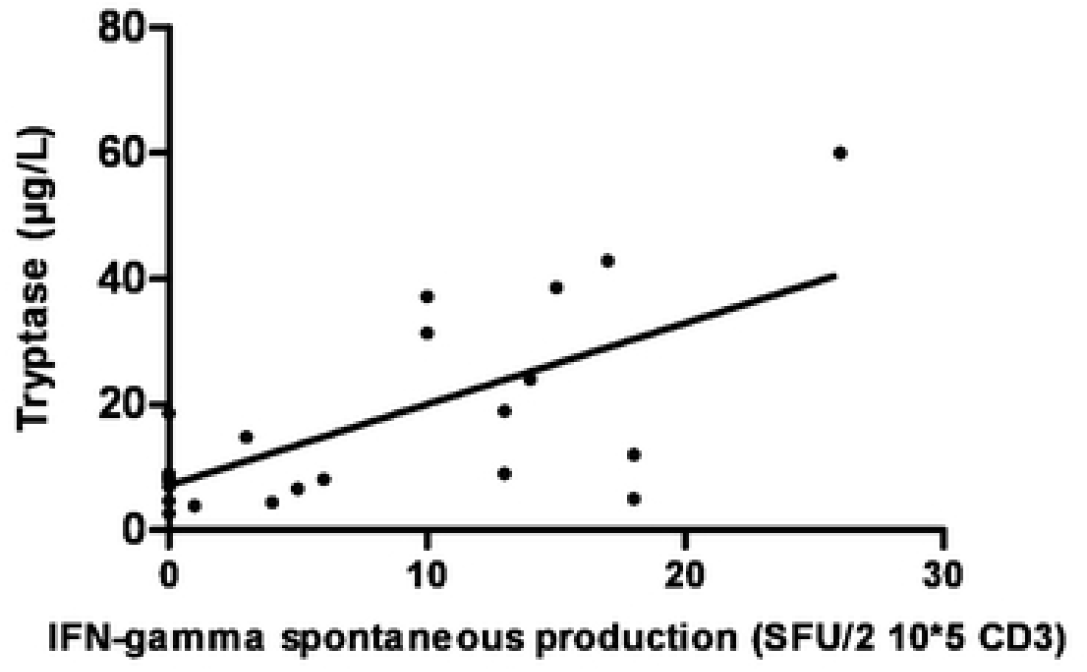
Correlation between basal tryptase level (μg/L) on Y axis and IFN-γ spontaneous production (SFU/2.10*5 CD3) on X axis observed on EliSpot assay. N=24 patients with CM, MIS and ISM. Linear regression: R^2^=0.44 (p<0.0004).

To our knowledge, this result has never been reported in the literature and may suggest some degree of additional protection against severe patterns of viral infections. Further works in our laboratory are currently performing to determine if this observation is related to a specific cytokine profile in patient plasma or due to a direct cellular mechanism between mast cells and T-cells. If confirmed, this specific phenotype in cMCADs patients might lead to therapeutic implications in the field of infectious diseases.

Overall, our results showed that cMCADs were able to develop effective and protective cellular and humoral response to SARS-CoV-2 but all of evaluable patients (4/4) with serial serology negated their serology after a median follow up of 33.0 weeks. Thus, anti-SARS-CoV-2 vaccination is strongly recommended, but its effectiveness remain to be confirmed in this specific population.

### Conclusion

In conclusion, non-advanced mastocytosis and monoclonal mast cells activation syndrome most likely do not confer an increased risk for severe COVID-19. A spontaneous IFN-γ production in patients with cMCADs may be involved in this observation and must be confirmed by further clinical and biological studies. If confirmed, this specific immune profile may explain protection against SARS-CoV-2 virus.

## Methods

### Patients

We have prospectively collected data from patients with cMCADs and COVID-19 documented by a positive SARS-CoV-2 PCR on nasal swab or with symptoms suggestive of COVID-19 associated with a positive anti-SARS-CoV-2 serology. Data were collected from the “Centre de référence des mastocytoses” (CEREMAST) rare diseases network in France. The study covered cases recorded from February 1st 2020 to February 1st 2021. cMCADs diagnosis were made according to WHO 2016 classification and clonal mast cells activation syndrome classification(1,23).

First, we sent a questionnaire to all patients over 18 years old with mastocytosis or MMAS with recent follow-up included in the CEREMAST national registry (N=828) and the “protocole physiopathologique de l’Association Française pour les Initiatives de Recherche sur le Mastocyte et les Mastocytoses (AFIRMM)”. The questionnaire sent collects the signs of mast cells activation displayed during the COVID-19, the current treatments and the specific signs and outcomes related to the COVID-19 presented by the patient. We then collected negative and positive cases for COVID-19 (Figure 1). Subsequently, we surveyed all competence centers in the French CERMAST network (N=24) to collect data from patients who had presented with COVID-19 but did not respond to the questionnaire. To ensure that there were no additional severe cases not reported, we made a request to the computerized registry (PMSI) at the Paris Public Hospitals Public Assistance (APHP) to search for possible cMCADs patients hospitalized for COVID-19 among the 8.3 million patients treated each year at the APHP. All patients with past COVID-19 performed an anti-SARS-CoV-2 serology.

### Ethic Statements

All patients with cMCADs were followed up in the CEREMAST network centers (mastocytosis reference centers in France). Patients were enrolled in a prospective, national, multicenter study sponsored by the French association for initiative and research on mast cell and mastocytosis (AFIRMM). This study was approved by the ethics committee of Necker Hospital and was carried out in compliance with the Declaration of Helsinki Principles protocol. A written informed consent was obtained (Comité de Protection des Personnes N°93-00). Blood samples were obtained as part of routine care in the follow-up for their cMCADs. Control cohorts were prospectively collected and analyzed as part of the COVID-HOP study (APHP200609).

### Immunological assays

The identification of SARS-CoV-2 specific T-cell responses was performed using an EliSpot that measure interferon-γ (IFN-γ) produced by specific SARS-CoV-2 T-cells. Briefly, Peripheral Blood Mononuclear Cells (PBMCs) were isolated from fresh blood collected during a follow-up consultation. After PBMC isolation by Ficoll density gradient, cells were stimulated for 18-20h using individual 15-mers 11-aa overlapping peptide pools of different SARS-CoV-2 proteins or common coronavirus proteins. Each responding cell was resulting in the development of one spot. Results were expressed as spot forming unit (SFU)/10^6^ CD3^+^ T-cells after subtraction of background values from wells with non-stimulated cells.

Negative controls were PBMC in culture medium (RPMI-1640, with L-glutamine and sodium bicarbonate (Sigma-Aldrich, Molsheim, France) supplemented with 10% human AB serum) without any stimulation. Positive controls were phytohemagglutinin PHA-P (Sigma-Aldrich) and CEFX Ultra SuperStim Pool (JPT Peptide Technologies GmbH, BioNTech AG, Berlin, Germany). SARS-Cov-2 peptide pools tested were derived from a peptide scan through SARS-CoV-2 Spike glycoprotein (2 pools: S1 for N-terminal fragment and S2 for C-terminal fragment), Membrane protein (M), Nucleoprotein (N), Envelope small membrane protein (E) and ORF3a protein. Pools of peptides derived from Spike glycoprotein of common Human alpha-coronavirus (HCoV-229E and HCoV-NL63) and beta-coronavirus (HCoV-OC43 and HCoV-HKU1) were also tested.

Humoral characterization, including anti-Spike SARS-CoV-2 IgG and IgA antibodies detection and neutralizing ability of anti-Spike IgG determination, was performed using previously described techniques: S-flow assay and S-pseudotype neutralization assays(24). Briefly, S-Flow assay used transduced Human embryonic kidney (HEK) 293T-cells encoding SARS-CoV-2 Spike protein. Cells were incubated with sera from patients (at a 1:300 dilution), and stained using either anti-IgG or anti-IgA. The fluorescent signal was measured by flow cytometry. For S-pseudotype neutralization assay, pseudotyped viruses carrying SARS-CoV-2 Spike protein were used. The viral pseudotypes were incubated with sera to be tested (at a 1:100 dilution), then added on transduced HEK 293T-cells expressing ACE2 and incubated for 48h at 37°c. The test measures the ability of anti-S antibodies to neutralize infection. Neutralization was calculated as described(24).

Control group including convalescent COVID-19 patients with mild to moderate and severe forms who were previously tested for SARS-CoV-2 EliSpots and serology in the Necker’s immunology laboratory.

### Statistics

Statistical analyses were performed using GraphPadPrism (version 6.0; GraphPad Software). Comparison tests were performed using Student’s t test, chi-square and Fisher’s exact tests when appropriate. The results are expressed as the mean or median +/- range [minimum; maximum]. P values < 0.05 were considered significant, values smaller than this are indicated in figure legends: *, P < 0.05; **, P < 0.01; ***, P < 0.001. ****, P < 0.0001.

## Abbreviations

APHP: Paris Public Hospitals Public Assistance
BMI: Body Mass Index
CEREMAST: Centre de Référence des Mastocytoses
CM: Cutaneous Mastocytosis
cMCADs: clonal Mast Cells Activation Disorders
IFN: Interferon
ISM: Indolent Systemic Mastocytosis
MIS: Mastocytosis in the Skin
MMAS: Monoclonal Mast cells Activation Syndrome
PBMC: Peripheral Blood Mononuclear Cells
SSM: Smoldering Systemic Mastocytosis
WHO: World Health Organization

## Acknowledgements

We would like to thank all the patients who responded to the questionnaire and the caregivers who took care of the patients with COVID-19.

## Author Contributions

### Conceptualization

J. Rossignol^1,2^, A. Ouedrani^3^, L. Chatenoud^3,4^ and O. Hermine^1,2†^

### Clinical data collection

J. Rossignol^1,2^, C. Bulai Livideanu^5^, S. Barete^6^, L. Terriou^7^, D. Launay^7^, R. Lemal^8^, C. Greco^1,2^, L. Frenzel^1,2^, C. Meni^1^, C. Bodemere-Skandalis^1,2^, L. Polivka^1,2^, A-F Collange^1,2^, H. Hachichi^1^, S. Bouzourine^1^, D. Nait Messaoud^1^, M. Negretto^5^, M. Gousseff^9^, S. Durupt^10^, J-C. Lega^10^, J-M Durand^11^, C. Gaudy^11^, G. Damaj^12^, M-P. Gourin^13^, M. Hamidou^14^, L. Bouillet^15^, E. Le Mouel^16^, A. Maria^17^, P. Zunic^18^, Q. Cabrera^18^, D. Vincent^19^, C. Lavigne^20^, E. Riviere^21^, C. Gourguechon^22^, A. Brignier^23^, L. Lhermitte^24^, T. J. Molina^2,25^, J. Bruneau^2,25^, J. Agopian^26^, P. Dubreuil^26^, D. Ranta^27^, A. Mania^8^, M. Arock^28^, O. Tournilhac^8^, O. Lortholary^1,2^, and O. Hermine^1,2†^

### Immunological assays (experiments and analysis)

A. Ouedrani^3^, L. Vendrame^3^, M.

Jambou^3^, I. Staropoli^29^, O. Schwartz^29^, L. Chatenoud^3,4^

### Supervision

O. Schwartz^29^, L. Chatenoud^3,4^ and O. Hermine^1,2†^

### Writing – original draft

All authors

## Supporting information

**Supplemental Figure 1 (S1):**
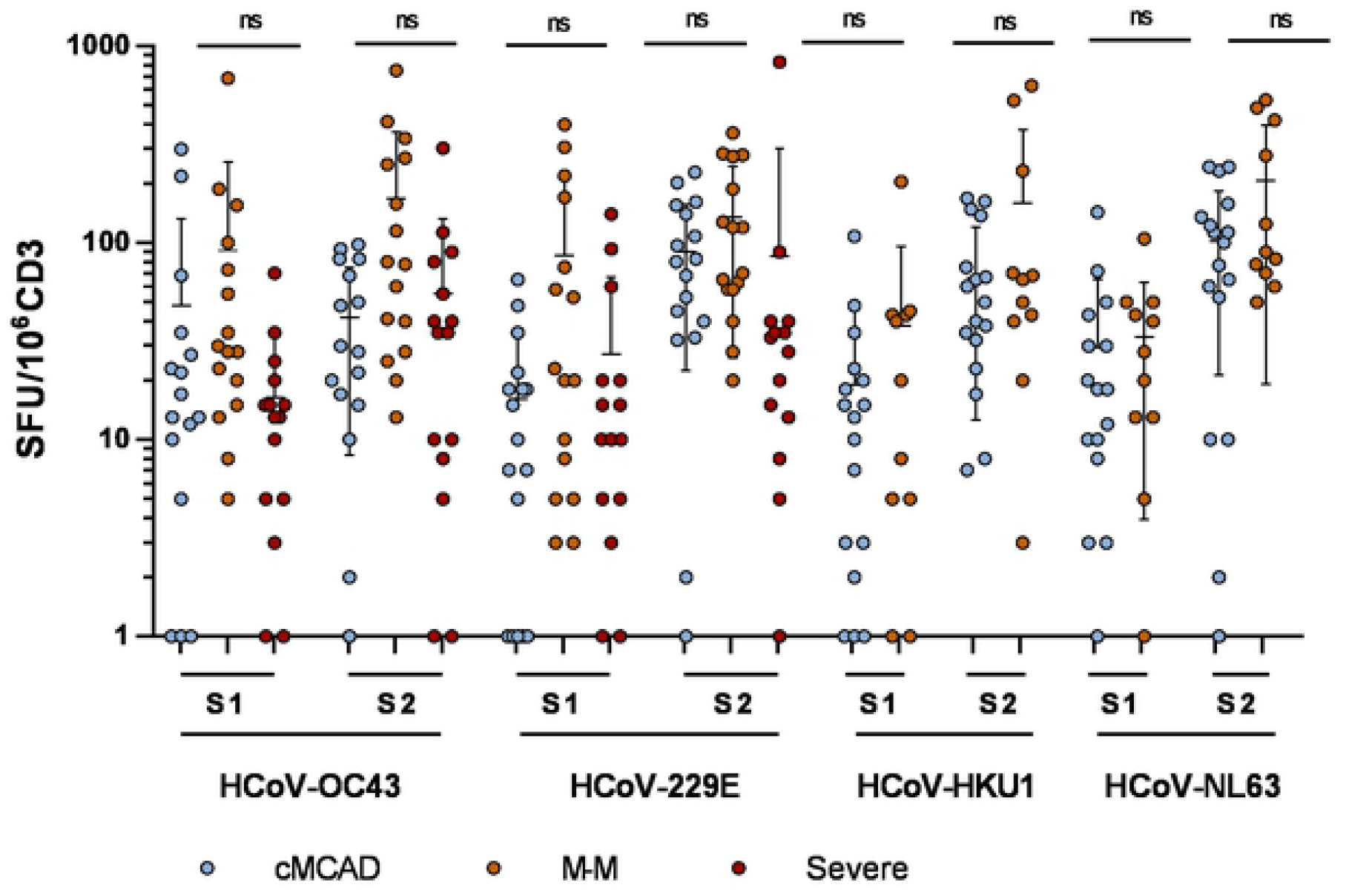
T-cells reactivities against common coronavirus in cMCADs, and no-cMCADs controls. Identification of HCoV-OC43, HCoV-229E, HCoV-HKU1 and HCoV-NL63 specific T-cells responses using EliSpot. Results were expressed as spot forming unit (SFU)/10^6^ CD3^+^ T-cells after subtraction of background values from wells with nonstimulated cells. Negative controls were patient cells in culture medium. Positive controls were peptides phytohemagglutinin PHA-P and CEFX Ultra SuperStim Pool. cMCADs: COVID-19 convalescent patients with clonal Mast Cells Activation Disorders. M-M: COVID-19 convalescent controls with mild or moderate COVID-19. Severe: COVID-19 convalescent controls with severe COVID-19. NS: non-significant.

**Supplemental Figure 2 (S2):**
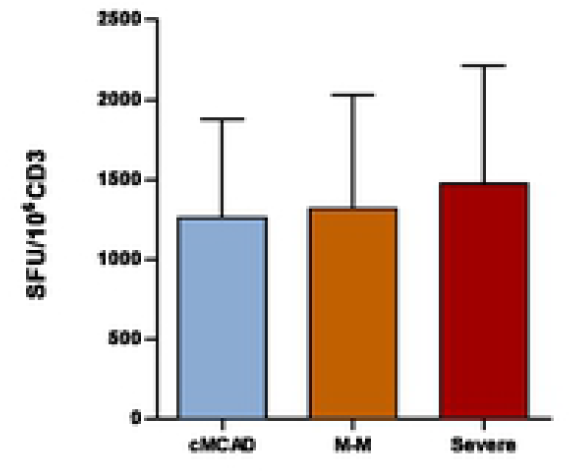
T-cells reactivities against CEFX Ultra SuperStim Pool. cMCADs: COVID-19 convalescent patients with clonal Mast Cells Activation Disorders. MM: COVID-19 convalescent controls with mild or moderate COVID-19. Severe: COVID-19 convalescent controls with severe COVID-19.

**Supplemental Figure 3 (S3):**
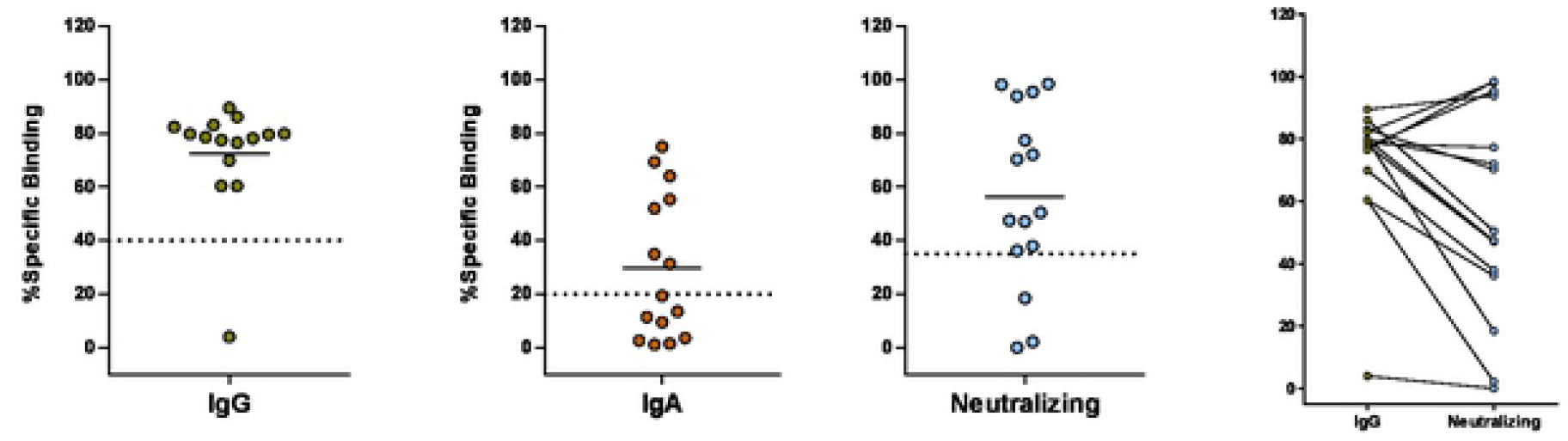
Two left panels: IgA and IgG serology determined with S-flow assay. The dashed line indicates the threshold of positivity. Third panel: Percentage IgG neutralizing ability determined with a viral pseudo-particle assay. Fourth panel: correspondence between the levels of anti-SARS-CoV-2 IgG antibodies and their neutralizing activity (Right).

